# WildAlert: A Real-Time, AI-Driven Early Warning System for Wildlife Health and Ecological Threat Detection

**DOI:** 10.64898/2026.04.07.716505

**Authors:** Pranav S. Pandit, Soumya Ranjan, Devin Dombrowski, Rachel Avilla, Carmen Ross, Deana L. Clifford, Krysta H. Rogers, Jane Riner, Heather Perry, Kirsten Gilardi, Martina Rutti, Leanne Flewelling, Katherine Hubbard, Terra Kelly

**Affiliations:** Department of Population Health and Reproduction, Weill School of Veterinary Medicine, University of California Davis, CA 95616, USA; Development Seed, Washington D.C., USA; The Wild Neighbors Database Project, Middletown, CA, USA; Wildlife Health Laboratory, California Department of Fish and Wildlife, Rancho Cordova, CA, 95670, USA; Wildlife Health Center, Weill School of Veterinary Medicine, University of California, Davis, Davis, CA, 95616, USA; Florida Fish and Wildlife Conservation Commission, Fish and Wildlife Research Institute, St Petersburg, 33701, Florida, USA; EpiEcos, Flagstaff, Arizona, USA

**Keywords:** Syndromic disease surveillance, Wildlife rehabilitation, Anomaly detection, Harmful algal bloom, Highly pathogenic avian influenza

## Abstract

Emerging infections and environmental disruptions increasingly threaten wildlife and ecosystem health. Free-ranging wildlife often serve as early indicators of ecological instability, making timely detection of morbidity and mortality events critical for early warning. Yet, existing systems lack the analytical capacity for real-time outbreak detection. We present WildAlert, an AI-driven early warning system that integrates fine-tuned BERT-based natural language processing models with unsupervised anomaly detection framework to identify unusual wildlife health events using real-time pre-diagnostic clinical data from wildlife rehabilitation organizations. The NLP module achieved high accuracy across clinical classifications and circumstances of admissions, enabling a pre-diagnostic syndromic surveillance framework. Retrospective validation demonstrated that WildAlert anomalies frequently coincided with or preceded confirmed morbidity events, including highly pathogenic avian influenza (HPAI), harmful algal bloom–associated toxicosis, cold-stunning in sea turtles, mass stranding events, West Nile virus, and mycoplasmosis. WildAlert establishes the world’s largest standardized, near real-time wildlife health surveillance system, transforming wildlife rehabilitation clinical records into actionable intelligence capable of detecting anomalies across taxa and regions, often before other surveillance methods. WildAlert provides a transferable analytical framework and scalable One Health model linking biodiversity monitoring, zoonotic disease preparedness, and ecosystem-linked environmental threats, with implications for conservation, public health and environmental hazard response.

## Introduction

Emerging infectious diseases and environmental hazards pose escalating threats to biodiversity, as free-ranging wildlife can serve as early indicators of ecological disruption and public health risk (Daszak et al., 2001; Johnson et al., 2020). These health threats, ranging from infectious pathogens to chemical toxicants, are increasingly recognized as significant drivers of wildlife morbidity and mortality. They often arise at the interface of ecological disruption and anthropogenic pressures, including habitat fragmentation, land-use change, climate shifts, and pollution. Integrated data streams encompassing human, wildlife, and livestock health information offer valuable opportunities for early disease detection and intelligence gathering on priority health threats. Participatory disease surveillance systems are increasingly employed to detect and respond to these threats at the human-animal-environmental interface (McNeil et al., 2022). However, these systems often exhibit substantial variation across data parameters and collection methodologies, hindering system interoperability and data sharing (McNeil et al., 2025). Passive syndromic surveillance approaches are also gaining traction for early disease detection. Passive syndromic approaches offer the ability to surveil for a wide range of diseases, including those caused by novel pathogens, and to detect health events prior to establishing definitive diagnoses (Lederberg et al., 2003), complementing more resource-intensive active surveillance.

Despite recent advancements in participatory and syndromic disease surveillance systems, their capacity for forecasting and early detection remains limited due to a lack of robust, scalable, and adaptable analytical algorithms. For example, while social media-based influenza-trackers in humans have achieved correlations of up to 89% with reported influenza activity (Achrekar et al., 2011), animal disease surveillance systems still rely predominantly on retrospective analyses (Kelly and Bland, 2006; King et al., 2023; Pruvot et al., 2023), limiting real-time responsiveness. Global livestock disease surveillance platforms like EMPRES-I, although capable of integrating near-real-time data streams, still lack advanced analytical pipelines necessary for proactive disease outbreak detection and forecasting (Claes et al., 2014). These gaps highlight the urgent need for improved algorithmic approaches that can enhance early warning capabilities, particularly in animal health surveillance.

As anthropogenic pressures intensify, so does the need for real-time, adaptive surveillance systems capable of detecting unusual wildlife health events at their onset (Ryser-Degiorgis, 2013; Sleeman et al., 2019). Wildlife rehabilitation organizations, which collectively admit millions of cases each year, represent a rich yet underutilized source of syndromic surveillance data. These organizations provide continuous, geographically broad access to species-level data and are well-positioned to detect early signals of disease outbreaks and other ecological disturbances (Cox-Witton et al., 2014; Kelly et al., 2021). Translating these data into actionable insights, however, requires robust analytical tools capable of processing unstructured clinical information, identifying meaningful patterns, and distinguishing true anomalies from background variation in a timely and scalable manner.

Recognizing this need, we developed WildAlert, a wildlife health surveillance platform that applies machine learning techniques to enhance the detection, classification, and interpretation of wildlife health events using real-time clinical data from wildlife rehabilitation organizations (Kelly et al., 2021). Currently deployed in California, Florida, Arizona, Washington, and Pennsylvania, the system has thus far relied on a relatively simple bag-of-words natural language processing (NLP) approach. This method does not account for word order or context, thereby resulting in a loss of semantic information and limiting the system’s ability to capture complex patterns in unstructured clinical text. In parallel, the platform’s time series analytical approach has been limited to moving averages of wildlife case admissions. This approach constrains the ability to accurately identify statistically significant anomalies that may represent true health events, particularly in the context of non-stationary and seasonally influenced data.

An improved current version of this system, WildAlert 2.0, addresses these limitations by incorporating a deep learning-based NLP large language model (LLM) built on bidirectional encoder representations from transformers (BERT). Unlike conventional approaches, this transformer-based model captures contextual relationships between terms, yielding improved classification accuracy (Sanh et al., 2020). In parallel, the baseline moving average-based anomaly detection method has been replaced with a more flexible, unsupervised anomaly detection framework incorporating Isolation Forest (IOF) and auto-encoder algorithms (AE), enabling enhanced detection of anomalies. By evaluating multivariate case data including taxonomy, temporal features, geography, and clinical classification, WildAlert 2.0 can identify both regional and statewide anomalies in wildlife admissions without reliance on manually defined baselines (Liu et al., 2012; Yu et al., 2024).

Building on recent advancements, the primary objective of this study is to rigorously evaluate the performance and representativeness of WildAlert 2.0 in detecting and characterizing unusual wildlife health events across taxa ranging from common to rare species. Specifically, we assess the utility of the integrated BERT-based natural language processing (NLP) models and two machine learning-based anomaly detection approaches in identifying wildlife morbidity and mortality events. To establish benchmarks for timeliness and accuracy of threat detection, we validate WildAlert-generated alerts against independently reported wildlife health data. In addressing limitations of existing wildlife health surveillance platforms, which often rely on retrospective analyses or lack advanced analytical pipelines, WildAlert 2.0 provides real-time, adaptive capabilities for early detection (Fig. 1). The platform’s geographical reach, active engagement of wildlife rehabilitation organizations, and demonstrated capacity to trigger targeted investigations underscore its potential as a reliable, scalable, and real-time surveillance tool. Taken together, these findings support the role of WildAlert 2.0 as both an early warning system for emerging ecological threats and a proactive mechanism for wildlife health monitoring.

**Fig. 1:**
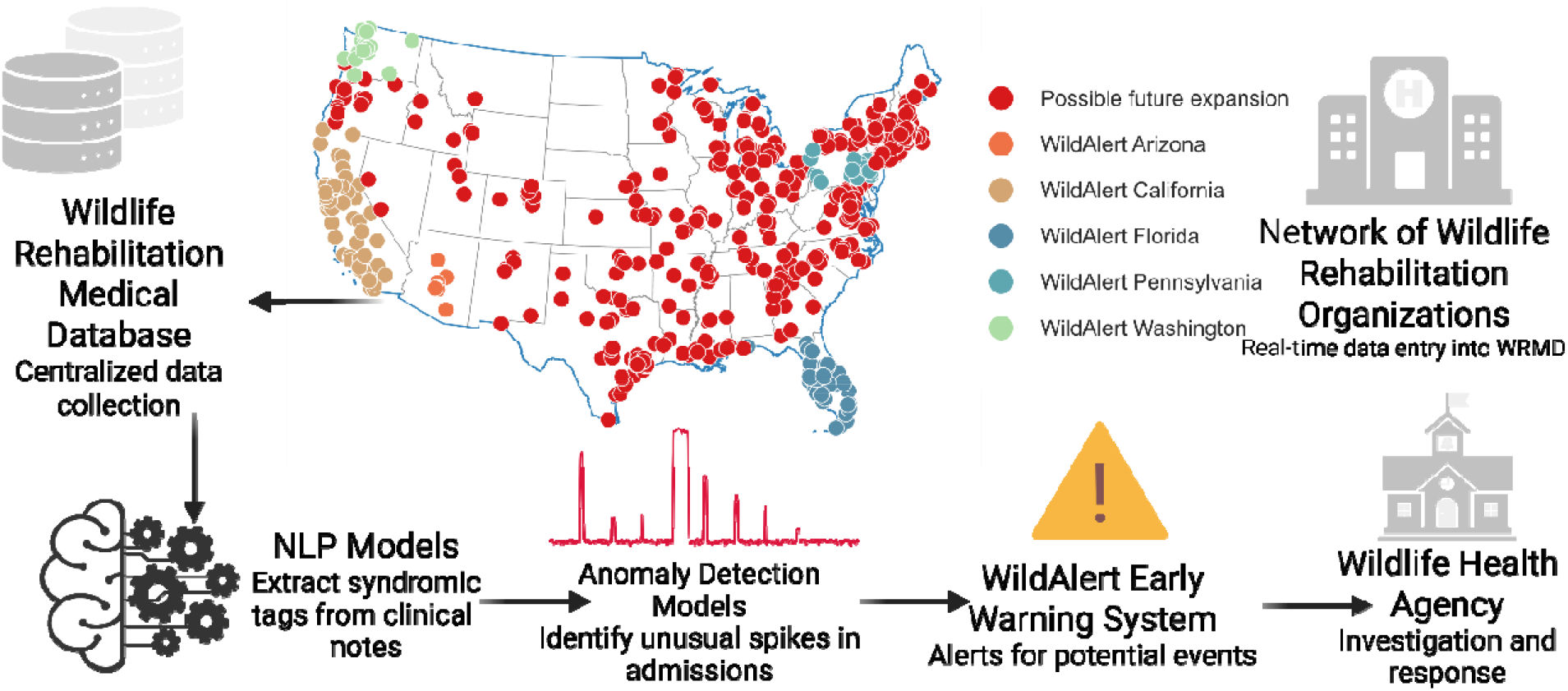
WildAlert Early Warning System for Wildlife Health Monitoring and Response. Figure illustrates the flow of real-time data from wildlife rehabilitation organizations across Arizona, California, Florida, Pennsylvania, and Washington into the Wildlife Rehabilitation Medical Database (WRMD). Using natural language processing (NLP) and anomaly detection models, WildAlert identifies syndromic patterns and temporal anomalies in wildlife admissions, enabling early detection of potential events and supporting timely investigation and response efforts by wildlife health agencies.

## Results

### Natural language processing (NLP) model performance

The fine-tuned BERT-based model demonstrated excellent performance in multi-label classification of clinical signs and causes of admission in rescued wildlife. Across 11 categories, the clinical classification model achieved a macro-averaged F1-score of 0.92, indicating strong overall performance and balance across both common and rare classifications (Supplementary Table 1). The micro-averaged F1-score was also 0.92, reflecting high overall accuracy. Class-specific F1-scores ranged from 0.88 to 0.97, with the highest performance in the Clinically Healthy (F1 = 0.97) and Physical Injury (F1 = 0.94) categories. Less frequent classes, such as Nonspecific (F1 = 0.88), were classified with robust predictive ability. To further evaluate classification behavior, we examined the confusion matrix shown in Fig. 2, which highlights the distribution of predicted versus true labels for the categories. Diagonal elements represent correct classifications, with accuracies ranging from 86% to 97%. The model was most accurate in identifying Clinically Healthy (97%), Physical Injury (94%), and Nutritional Disease (92%) cases. Systematic misclassifications were observed, including 5% of Dermatologic Disease cases misclassified as Urogenital Disease, and 5% of Nonspecific cases misclassified as Clinically Healthy, suggesting overlapping clinical sign descriptions or annotation ambiguity. Occasional misclassification occurred for Ocular Disease and Neurologic Disease cases, which were occasionally labeled as Urogenital Disease or Physical Injury, respectively. A small proportion of cases across multiple classes (e.g., Respiratory Disease, Hematologic Disease, Nutritional Disease) were not assigned to any label, indicating areas where the model may benefit from improved handling of ambiguous or underrepresented text patterns.

**Table 1:** Real-world wildlife health events detected by WildAlert and independently confirmed. Select unusual wildlife health event alerts generated by WildAlert in Florida and California (2024-2025). The table includes date, affected taxa, clinical classifications, geographic locations, and confirmed event diagnoses. *Entries without a clinical classification represent all admissions for the given taxa, regardless of specific clinical signs.

| State | Date of Alert | Species/Taxa | Clinical Classification(s) | Geographic Location(s) | Event Diagnosis |
| --- | --- | --- | --- | --- | --- |
| Florida | 6/17/24 | Greater Shearwaters |  | East/Atlantic Coast of FL | Stranding due to adverse weather |
|  | 9/3/24 | Seabirds | Neurologic Disease | Gulf | Brevetoxicosis |
|  | 9/3/24 | Shorebirds | Neurologic Disease | Gulf | Brevetoxicosis |
|  | 1/18/25 | Sea turtles |  | Gulf Coast | Cold stunning |
| California | 4/25/24 | Brown Pelicans | Nutritional Disease | Central and South Coast | Starvation |
|  | 1/25/25 | Sanderlings | Neurologic Disease | Central CA Coast | Highly Pathogenic Avian Influenza |
|  | 2/27/25 | Mesocarnivores | Neurologic Disease, Dermatologic Disease, and Ocular Disease | Central and Southern California | canine distemper virus |
|  | 3/23/25 | Seabirds | Neurologic Disease and Nutritional Disease | Southern CA | Domoic acid toxicosis |

**Fig. 2:**
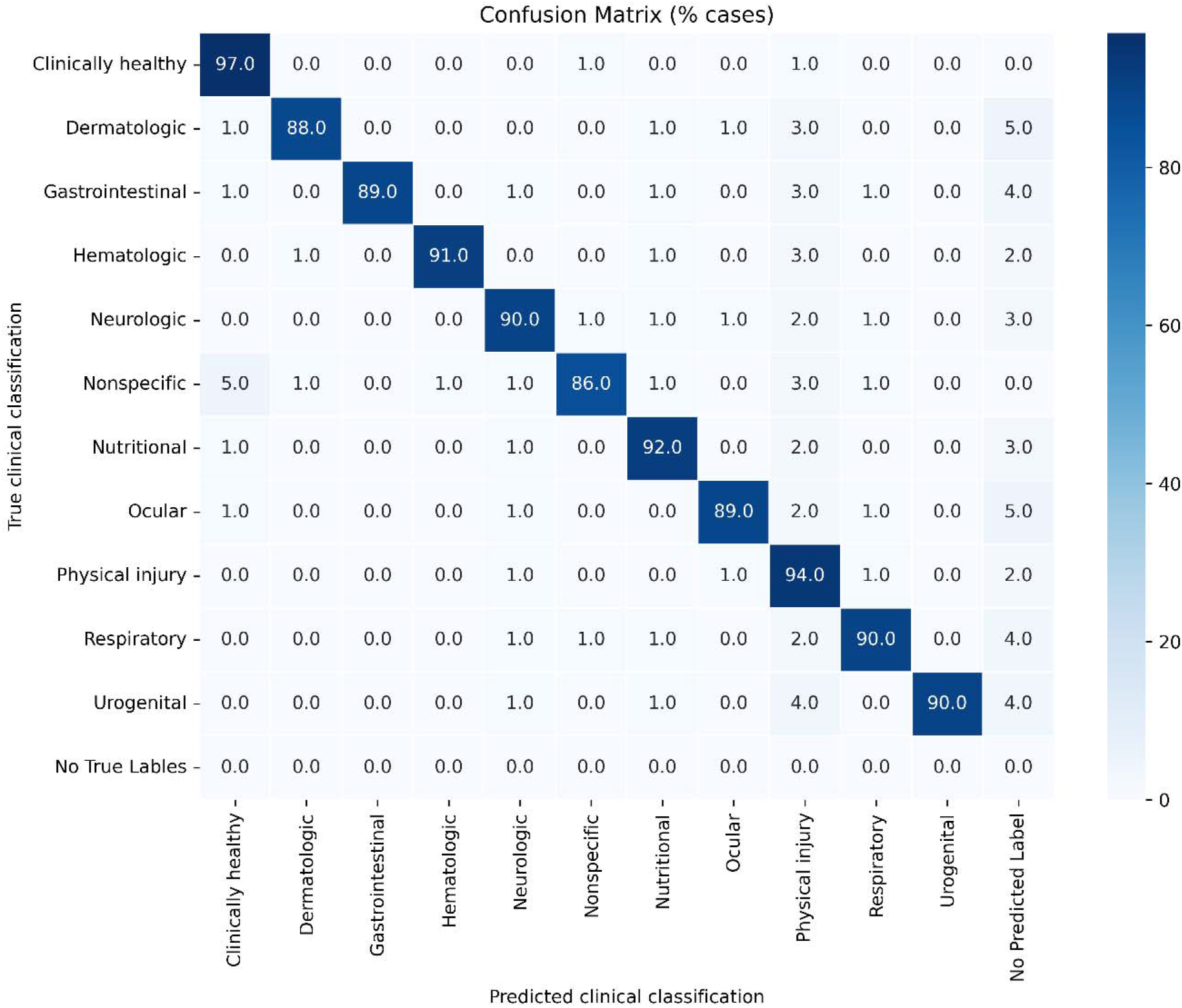
Performance of the NLP model for clinical classification across wildlife cases. The figure presents the confusion matrix for the model’s predictions across 11 clinical categories. Each cell represents the percentage of cases predicted for a given class (columns) relative to their true class (rows). The diagonal elements indicate correct classifications, while off-diagonal elements represent misclassifications.

For circumstances of admission, the model demonstrated high discriminative performance in classifying 74 distinct categories based on free-text narratives (Supplementary Table 2). Precision and recall exceeded 0.95 for most categories, with particularly strong performance for well-defined circumstances such as gunshot, and plane collision (F1 = 1.000), as well as common circumstances like cat interaction (F1 = 0.989) and vehicle collision (F1 = 0.986). The confusion matrix (Supplementary Fig 1) demonstrated high classification accuracy across many frequently observed circumstances for admission. However, performance declined for rare or semantically ambiguous classes.

### Anomaly detection model performance

Tagging of wildlife rehabilitation cases with clinical classifications and circumstances of admissions generated *WildAlert targets* representing a distinct surveillance time series characterized by species-syndrome-location to be monitored for anomalous activity. To evaluate the robustness and sensitivity of machine learning-based anomaly detection models under realistic surveillance conditions, we conducted a comprehensive sensitivity analysis across 32 surveillance WildAlert targets representing both common (n= 17, Supplementary Fig 1) and rare species (n= 15, Supplementary Fig 2). These were based on baseline reporting frequency. Endemic targets exhibited sustained, recurrent baseline activity throughout the study period, whereas sporadic targets showed infrequent or intermittent reports separated by extended zero-count intervals. We systematically perturbed the input time series with stochastic outliers and gradual linear trends to mimic real-world noise and shifting baselines and assessed model performance across a range of sliding window sizes (3–20 weeks) and contamination levels (anomaly injection rates, 1–20%) to capture variation in temporal resolution and anomaly prevalence.

We evaluated inter-model agreement and consistency in anomaly detection using multiple metrics. For common endemic WildAlert targets, the overlap in anomalies detected by both the IOF and AE models was 19.3% (SD = 8.98%) with a Cohen’s Kappa of 0.25 (SD = 0.12), indicating fair agreement. The Jaccard similarity coefficient averaged 0.16 (SD = 0.08), reflecting limited overlap among flagged events. The autoencoder model showed a more balanced sensitivity profile, with moderate sensitivity to contamination level (β =- 1.3), smaller effects from trend (β = 0.13), outliers (β = –2.42), and window size (β = –0.15) (Fig. 3a). In contrast, the Isolation Forest model exhibited extreme sensitivity to contamination level (β = 282.37), moderate sensitivity to outlier injection (β = 3.02), and minimal influence from trend (β = -0.04) and window size (β = 0.10) (Fig. 3b).

**Fig. 3:**
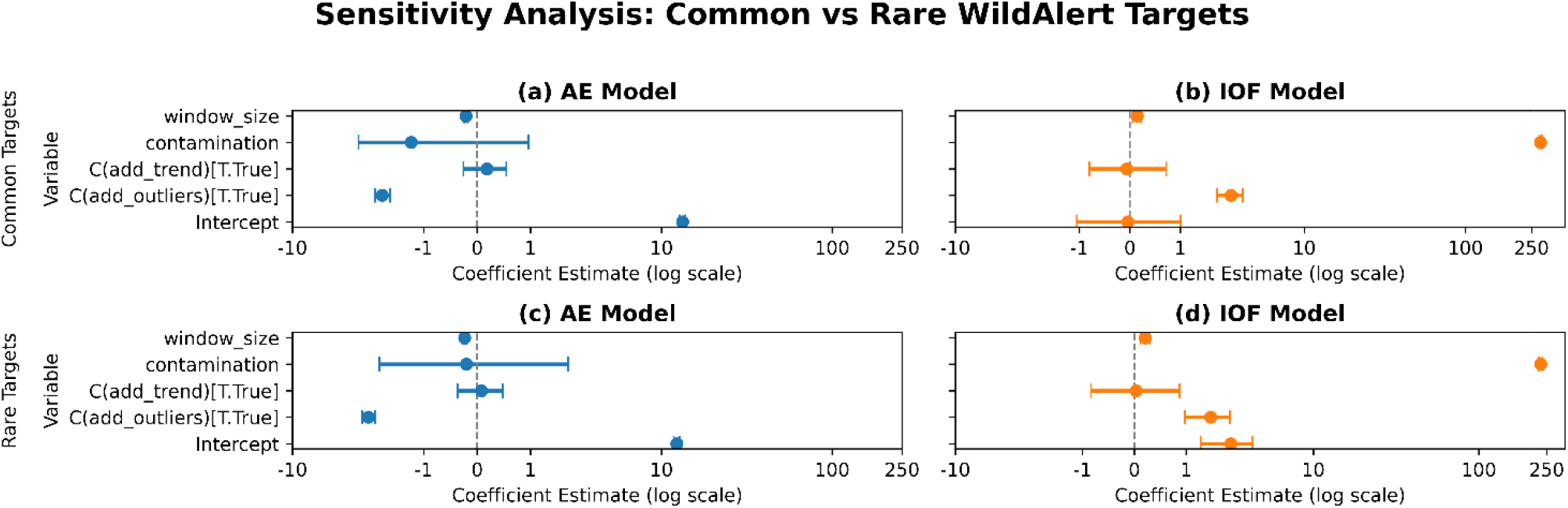
Sensitivity of anomaly detection models to perturbations in surveillance data. Comparison of anomaly detection model sensitivity to data perturbation parameters using standardized regression coefficients shown for four predictors (contamination level, outlier injection, trend strength, and window size) across two anomaly detection models: Autoencoder (left) and Isolation Forest (right). The x-axis represents the magnitude and direction of each coefficient on a logarithmic scale, indicating the relative influence of each parameter on the number of anomalies detected. Positive values denote increased anomaly detection with higher parameter values, while negative values indicate suppression.

For rare (sporadic) WildAlerts targets, the overlap in anomalies detected by both the Isolation Forest and Autoencoder models was 18.8% (SD = 10.61%) with Cohen’s Kappa between the two models was 0.24 (SD = 0.13), also indicating fair agreement. The Jaccard similarity coefficient averaged 0.15 (SD = 0.08), reflecting limited overlap among flagged events. Similar to analysis done for common (endemic) WildAlert targets, the AE model showed high sensitivity to trend (β = –3.10), with moderate sensitivity to contamination level (β =- 0.14), and smaller effects for trend (β = 0.05) on the number of detected anomalies (Fig. 3c), and the isolation forest model also exhibited extreme sensitivity to contamination level (β = 230.12), moderate sensitivity to outlier injection (β = 1.78) and minimal influence from trend (β = 0.02) and window size (β = 0.15) (Fig. 3d).

### Detection of HPAI-associated wildlife health events

WildAlert demonstrated variable performance in detecting wildlife health anomalies temporally aligned with USDA-confirmed HPAI detections across multiple avian taxonomic groups and states. In California, USDA reported HPAI in raptors across four distinct multi-month intervals: July 2022-April 2023, July-August 2023, October 2023-April 2024, and November 2024-March 2025 (Fig. 4A). During the first interval (July 2022-April 2023), the IOF model detected multiple anomalies, and the AE flagged a single anomaly. In the second and third intervals, the IOF model detected a single anomaly in each interval, while the AE model detected no anomalies. Notably, during the fourth interval, both models detected multiple overlapping anomalies with high signal intensity. Statistical comparisons of anomaly intensity scores across all four intervals, comparing USDA-confirmed HPAI periods to outside those intervals, yielded significant results for the AE model (t-test p = 0.028; permutation p = 0.007) and non-significant results for the IOF model (t-test p = 0.195; permutation p = 0.142, for IOF model). For California shorebirds with neurologic disease, multiple USDA-confirmed HPAI detections occurred between December 2022 and February 2025. Both AE and IOF models flagged anomalies during several of these intervals. However, statistical comparisons of anomaly intensity scores during USDA-confirmed periods to outside those intervals yielded non-significant results for both models (AE: t-test p = 0.17, permutation p = 0.37; IOF: t-test p = 0.24, permutation p = 0.36), suggesting elevated but not statistically distinct signal activity (Fig. 4B). Additionally, a Sanderling (*Calidris alba*) stranding event investigated by CDFW in January 2025 that was determined to be caused by HPAI coincided with an anomaly detected by the WildAlert IOF model.

**Fig. 4:**
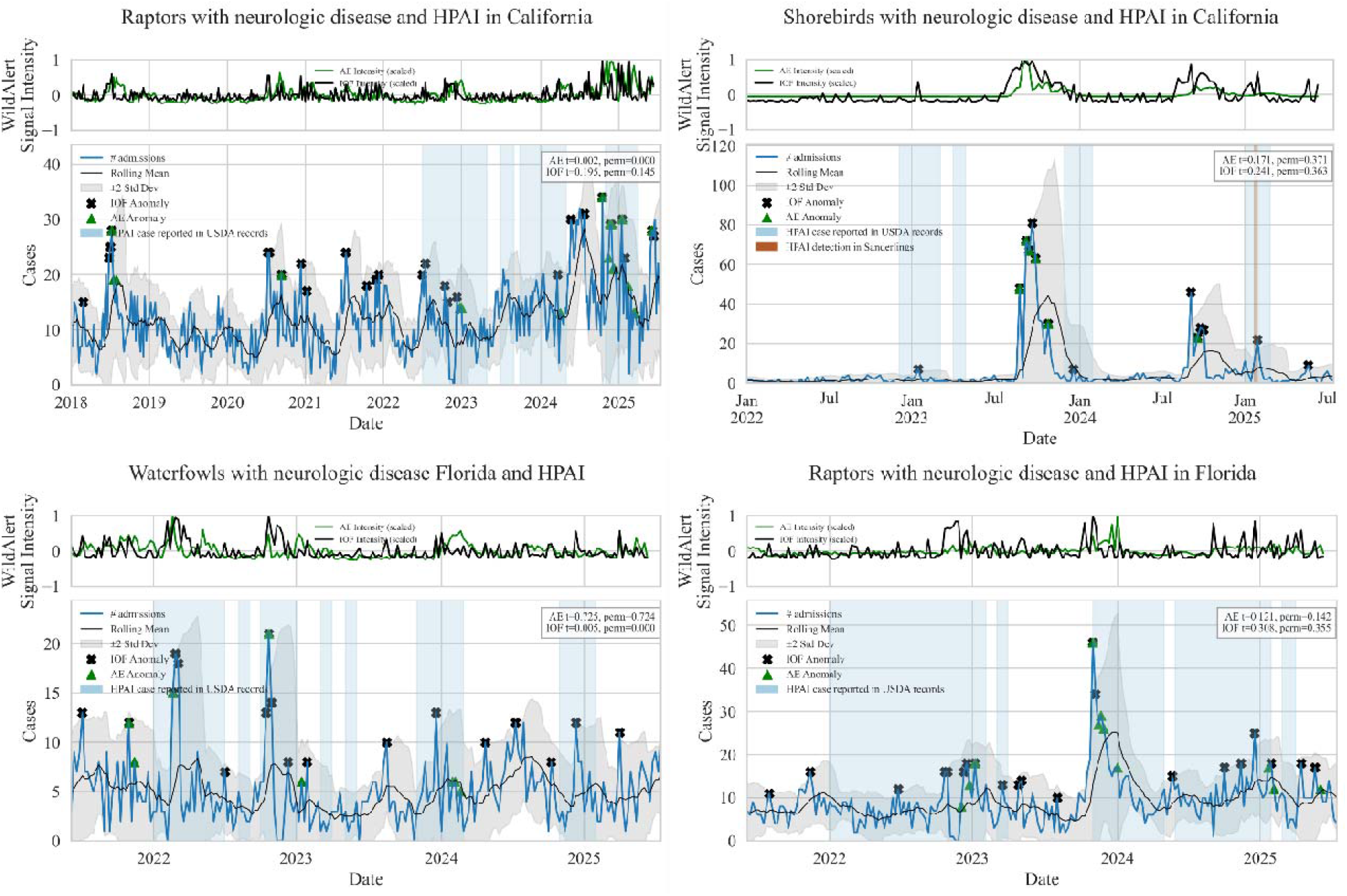
Detection of HPAI-associated anomalies by WildAlert across regions and taxa. WildAlert anomaly signals for neurologic disease in raptors, shorebirds, and waterfowl across California and Florida, overlaid with USDA-confirmed HPAI detection periods (shaded regions). Each panel shows weekly case counts and anomaly intensities from Autoencoder (AE) and Isolation Forest (IOF) models. Green triangles and black crosses denote anoimes detected through autoencoder and isolation forest models, respectively. AE and IOF signal intensities are plotted on the secondary y-axis.

Consistent HPAI detections in Florida waterfowl spanned seven distinct intervals from January 2022 to January 2025. WildAlert detected anomalies during several of these periods. Notably, the IOF model showed statistically significant signal elevation during USDA-confirmed periods (t-test p = 0.0048; permutation p < 0.001), while the Autoencoder model did not (t-test p = 0.73; permutation p = 0.72), indicating greater sensitivity of the IOF model to HPAI-associated disruptions in this WildAlert target (Fig. 4C). A surveillance target of raptors with neurologic disease in Florida led to anomalies overlapping across five confirmed HPAI detections between January 2022 and March 2025. However, statistical comparisons yielded non-significant results for both models (AE: t-test p = 0.12, permutation p = 0.14; IOF: t-test p = 0.31, permutation p = 0.36).

### Wildlife morbidity and mortality events detected by WildAlert (2024–2025)

In 2024 and 2025, WildAlert played a pivotal role in detecting multiple wildlife health events across Florida and California (Table 1), with detections cross-validated against independent monitoring programs and laboratory testing conducted by state wildlife agencies and wildlife rehabilitation organizations. In mid-June 2024, the system flagged an unusual stranding event involving Greater Shearwaters (*Ardenna gravis*) along Florida’s Atlantic Coast, later attributed to adverse weather conditions by the Florida Fish and Wildlife Conservation Commission (FWC). In September and October 2024, neurologic disease alerts were triggered for seabirds and shorebirds, which included Laughing Gulls (*Leucophaeus atricilla*) and Ruddy Turnstones (*Arenaria interpres*), along Florida’s Gulf Coast. Subsequent investigations by the FWC in collaboration with wildlife rehabilitation organizations confirmed brevetoxicosis resulting from exposure to *Karenia brevis*-derived neurotoxins (brevetoxins) as a contributing factor to the stranding event (Fig. 5-6). In January 2025, an alert was triggered for sea turtles along Florida’s Gulf Coast following a very large-scale cold-stunning event that affected over 1,000 green sea turtles (*Chelonia mydas*), with fewer numbers of loggerhead (*Caretta caretta*) and Kemp’s ridley (*Lepidochelys kempii*) turtles also impacted.

**Fig. 5:**
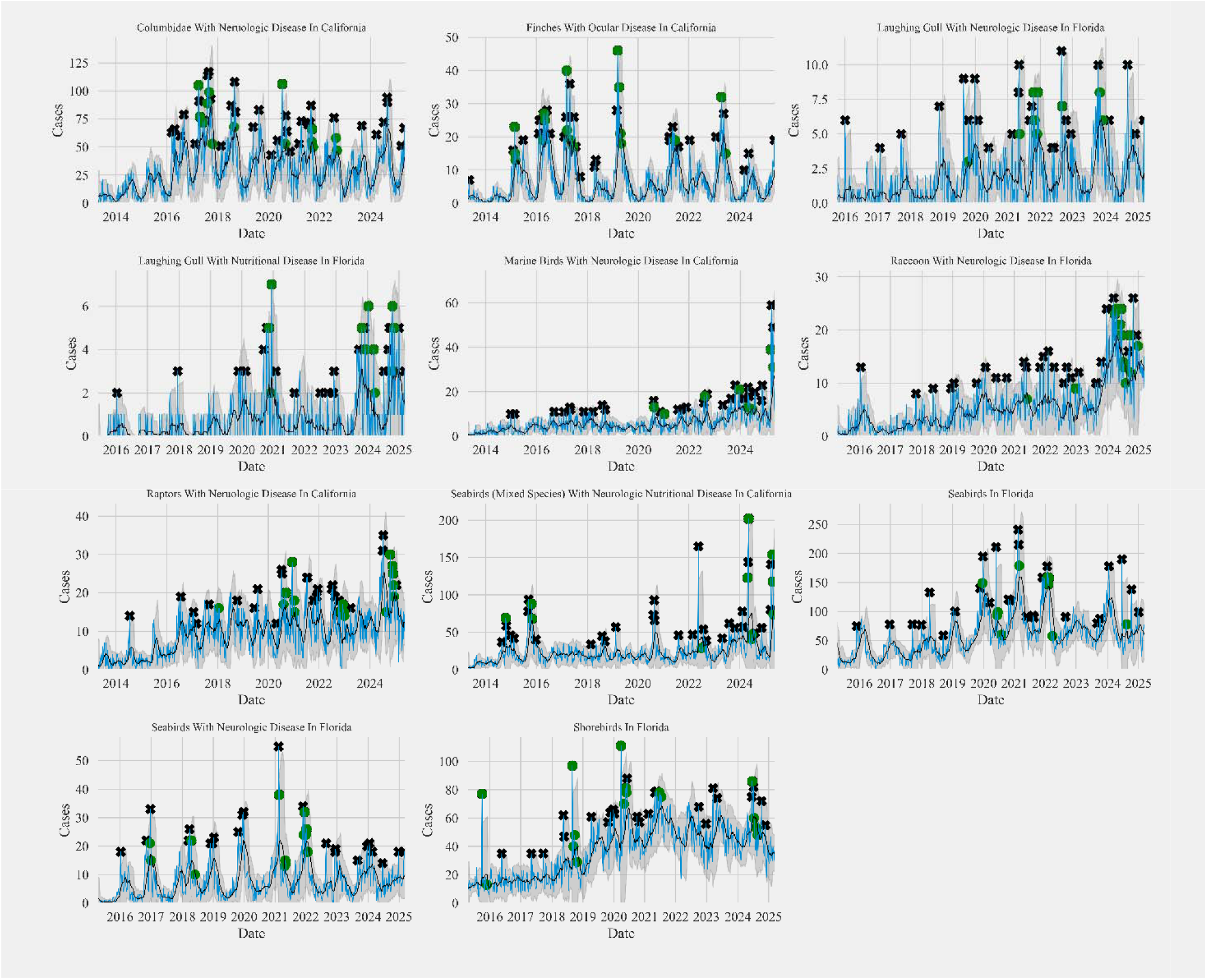
Time series of wildlife health signals illustrating diverse anomaly detection patterns in commonly found taxa-clinical classification targets. Subplots show: (a) Columbids with neurological disease in California, (b) Finches with ocular disease in California (c) Laughing Gulls with neurological disease in Florida, (d) Laughing Gulls with nutritional disease in Florida, (e) Marine birds with neurological disease in California (f) Raccoons with neurologic disease in Florida, (g) Raptors with neurological disease in California, (h) Seabirds with neurological and nutritional disease in California, (i) All seabirds admissions in Florida, (j) Seabirds with neurologic disease in Florida, and (k) All shorebird admissions in Florida. Blue lines represent monthly admissions, black lines indicate the rolling mean, and gray shaded areas denote the

**Fig. 6:**
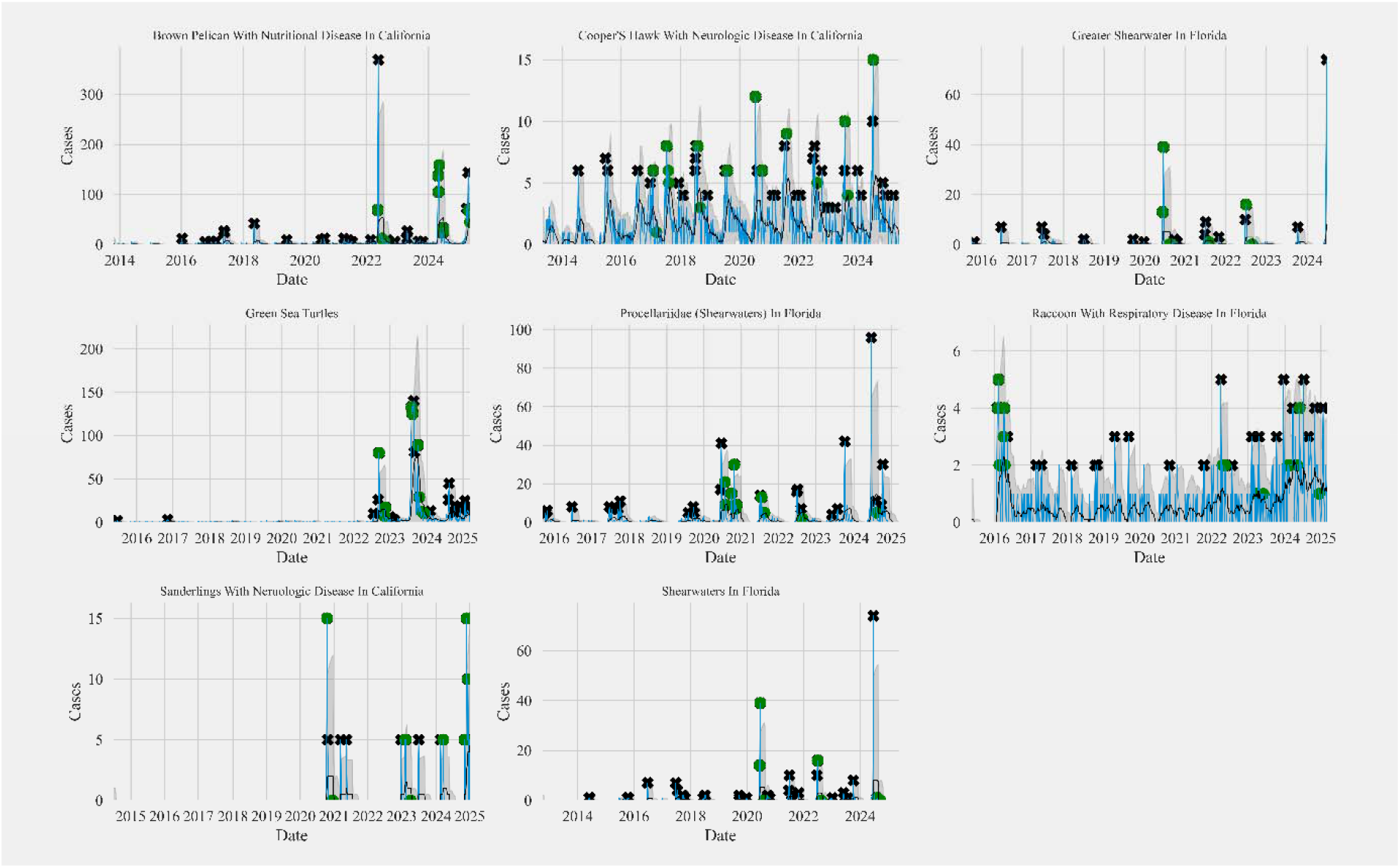
Time series of wildlife health signals illustrating diverse anomaly detection patterns in rare taxa-clinical classification targets. Subplots show: (a) Brown Pelicans with Nutritional Disease in California, (b) Cooper’s Hawk with Neurological disease in California, (c) All Greater Shearwaters admissions in Florida, (d) All Green sea turtles admissions in Florida, (e) All Procellariidae (Shearwaters) admissions in Florida, and (f) Raccoons with respiratory disease in Florida (g) Sanderlings with neurological disease in California (h) All Shearwater species admissions in Florida. Blue lines represent monthly admissions, black lines indicate the rolling mean, and gray shaded areas show the rolling standard deviation. Anomalies detected by the Isolation Forest model are marked with black stars, and anomalies identified by the Autoencoder model are shown as green circles.

In California, WildAlert detected a significant nutritional disease event in Brown Pelicans (*Pelecanus occidentalis*) beginning in April 2024, spanning the coast from Monterey to Orange County (Duerr et al., 2025). Post-mortem examinations revealed starvation, with many birds also having injuries related to fishing gear. The primary cause of the stranding event was reported to be a lack of adequate food resources. In January 2025, WildAlert flagged Sanderlings admitted to wildlife rehabilitation organizations along the central coast with neurologic disease, with subsequent post-mortem examinations and diagnostic testing confirming highly pathogenic avian influenza (HPAI) in several individuals. In February 2025, an alert was triggered for mesocarnivores in central and southern California, including striped skunks (*Mephitis mephitis*), raccoons (*Procyon lotor*), and gray foxes (*Urocyon cinereoargenteus*), presenting with neurologic, dermatologic, and ocular disease. The pattern of clinical signs was consistent with canine distemper virus in these animals and testing confirmed the diagnosis in a number of these cases. A major multi-species event was detected in March 2025 in southern California, involving Brown Pelicans, Western Grebes (*Aechmophorus occidentalis*), Brandt’s Cormorants (*Urile penicillatus*), and Red-throated Loons (*Gavia stellata*). Affected birds exhibited a combination of neurologic and nutritional disease signs and post-mortem and diagnostic testing confirmed domoic acid toxicosis (Fig. 5-6).

### Cross-correlation of marine bird admissions with Karenia brevis bloom activity

The cross-correlation analysis of *Karenia brevis* counts and WildAlert-reported marine bird admissions in Florida (Fig. 7) revealed species-specific temporal patterns associated with foraging ecology, with species clustering into rapid, intermediate, and delayed response groups relative to HAB conditions, reflecting potential differences in timing of exposure or response to bloom events. Pelagic and near-surface foragers exhibited the strongest and most temporally proximal associations. Sooty Terns (*Onychoprion fuscatus*), showed a strong positive correlation at lag +6 (r = 0.606) and lag +1 (r = 0.433), both exceeding the confidence interval (±0.377), indicating probable direct exposure to brevetoxin.

**Fig. 7:**
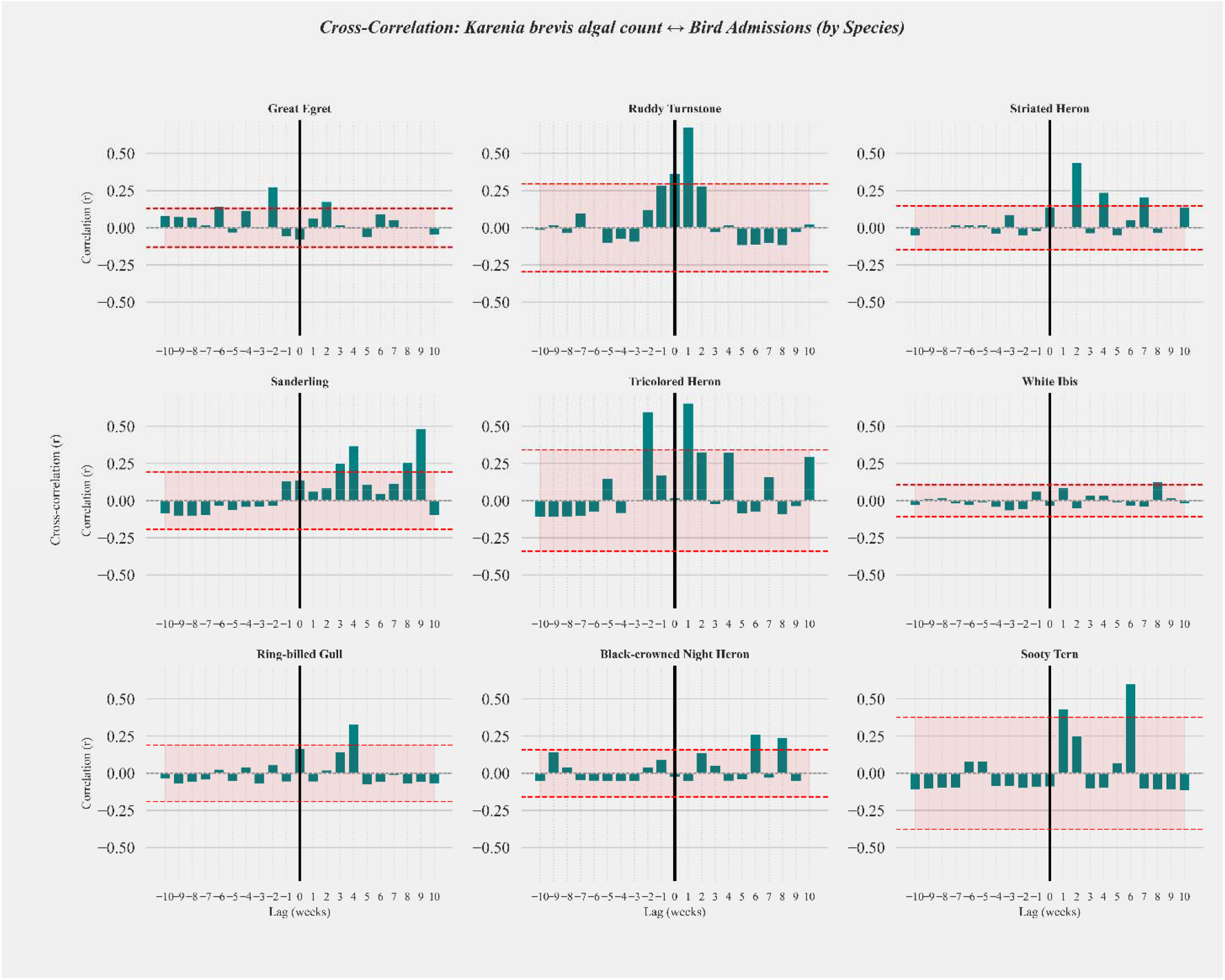
Species-specific temporal associations between bird admissions and harmful algal bloom activity. Species-specific cross-correlation coefficients (r) between standardized weekly admissions reported by WildAlert Florida and mean weekly Karenia brevis algal counts across lags from –10 to +10 weeks. Positive lags indicate bird admissions preceding HAB activity, while negative lags reflect HAB activity preceding bird admissions. Red dashed lines represent Bartlett-adjusted 95% confidence intervals.

Ruddy Turnstones (*Arenaria interpres*), which forage in the intertidal zone, also exhibited a strong positive correlation at lag +1 week (r = 0.679, 95% CI ±0.296), with additional elevated correlations at lag 0 (r = 0.365) and lag –1 (r = 0.289). These patterns suggest close temporal alignment between HAB activity and strandings in this species, consistent with acute exposure or rapid ecological response.

Species that forage primarily on benthic prey or exploit a broader range of food sources showed more delayed and diffuse responses. Sanderlings (*Calidris alba*) demonstrated a gradual increase in positive correlation from lag +3 to +9 weeks, peaking at lag +9 (r = 0.485, 95% CI ±0.194). This extended lag structure suggests that strandings may reflect cumulative or indirect effects of HABs, such as trophic disruption or migratory stress. Ring-billed Gull (*Larus delawarensis*), an opportunistic omnivore, showed a notable positive correlation at lag +4 (r = 0.331, 95% CI ±0.190), with additional peaks at lag 0 (r = 0.169) and lag +3 (r = 0.144). These results suggest a moderately delayed response to HAB conditions.

Wading birds using estuarine and nearshore habitats displayed heterogeneous lag patterns. Green Herons (*Butorides virescens*) showed a peak correlation at lag +2 weeks (r = 0.440, 95% CI ±0.148), with secondary peaks at lag +4 (r = 0.239) and lag +7 (r = 0.207). These findings suggest a moderately delayed response to HAB conditions, possibly mediated by habitat-level changes or prey availability. Tricolored Heron (*Egretta tricolor*) showed a sharp spike in correlation at lag –2 (r = 0.599) and lag +1 (r = 0.658), both exceeding the confidence interval (±0.342). These dual peaks suggest both anticipatory and reactive relationships with HABs, possibly reflecting complex behavioral or physiological responses. Black-crowned Night Herons (*Nycticorax nycticorax*) displayed elevated correlations at lag +6 (r = 0.262) and lag +8 (r = 0.242, 95% CI±0.159). These patterns suggest a delayed but potentially meaningful association with HAB intensity. In contrast, species less tightly coupled to marine food webs showed weaker associations. White Ibis (*Eudocimus albus*) exhibited relatively weak correlations across all lags, with a modest peak at lag +8 (r = 0.127). Overall, the analysis revealed that positive correlations between strandings and HAB activity were most pronounced at short positive lags (1–4 weeks) for several species, suggesting that HABs may act as proximal drivers of strandings. In contrast, negative correlations at earlier lags (–8 to –1 weeks) were observed in some species, suggesting that strandings may also precede or coincide with bloom development, potentially serving as early ecological indicators.

## Discussion

Our findings demonstrate that WildAlert 2.0, an AI-enhanced wildlife health surveillance system, effectively integrates natural language processing and anomaly detection algorithms to identify unusual patterns in clinical wildlife data in near real-time. Traditional approaches to early warning systems for wildlife disease have relied on manual review and basic statistical trend detection, limiting scalability and timeliness. By leveraging deep learning and unsupervised machine learning approaches, this platform represents a significant advancement in monitoring, enabling early detection of morbidity and mortality events across a wide range of taxa, including both common and rare species, as well as infectious and noninfectious threats.

The BERT-based NLP models in WildAlert exhibited strong performance in automatically classifying clinical presentations and circumstances of admission from unstructured wildlife rehabilitation records. This automated classification enables rapid structuring of complex case data, reducing reliance on manual coding and improving data usability for downstream analytics. Performance was robust across multiple clinical and situational categories, including those with imbalanced representation, suggesting that advanced language models can generalize well in the wildlife health context despite the inherent diversity and complexity of such data. These results support the use of large language models for extracting structured, biologically meaningful information from wildlife health records, a critical step towards scalable syndromic surveillance. Comparative sensitivity analysis of the two anomaly detection models revealed key differences in their robustness and parameter stability under conditions mimicking real-world surveillance data streams. The IOF model was highly sensitive to the anomaly injection rate (contamination parameter), indicating that its output is heavily influenced by assumptions about the proportion of anomalous data. While moderately responsive to injected outliers, IOF exhibited minimal sensitivity to underlying temporal trends and temporal window size, suggesting it performs well at detecting abrupt and discrete anomalies, but is less effective in capturing gradual shifts and may be prone to overfitting when parameterized for high sensitivity. In contrast, the AE model showed a more balanced response profile, with moderate sensitivity to contamination, slight negative influence from trend and outliers, and a pronounced negative effect of increasing temporal window size. AE’s stable anomaly counts across perturbed datasets highlight its robustness to non-stationary signals and background noise. These findings align with prior evaluations showing that autoencoders in health surveillance, are more resilient to gradual baseline drift and well-suited for early detection in complex datasets with seasonality (Wen et al., 2021). Taken together, these results support a hybrid approach, leveraging both models’ strengths: AE for robustness to evolving baselines and IOF for sensitivity in detecting abrupt, high-impact events. The moderate inter-method agreement further suggests that combining both methods in an ensemble enhances detection of diverse anomaly types (Sun, 2011).

WildAlert successfully detected a range of unusual wildlife health events, subsequently confirmed through investigations by state wildlife agencies. These included outbreaks of known pathogens (e.g., highly pathogenic avian influenza, canine distemper virus), as well as non-infectious events (e.g., HAB-associated toxicity, cold-stress in sea turtles, and marine bird strandings due to HAB-associated toxicities and extreme weather). These diverse outcomes illustrate WildAlert’s flexibility in identifying both infectious and non-infectious health threats in near real-time, reinforcing its value for agencies tasked with surveillance and event response. Cross-correlation analysis revealed that strandings of Ruddy Turnstones and Tricolored Herons typically occurred about 1–2 weeks after peak Karenia brevis abundance, while Sooty Tern strandings lagged by approximately 2–3 weeks, indicating species-specific temporal responses to bloom events. WildAlert 2.0 anomaly alerts frequently coincided with or preceded these events, demonstrating the system’s capacity to detect early indicators of environmental disruption. This ability to temporally align wildlife health data with known ecological hazards is critical for timely intervention and resource allocation. Beyond early detection, WildAlert also provides insight into the true scale of wildlife morbidity and mortality events, in otherwise substantial underestimated impact due to limited human encounters of wildlife and much less testing in other surveillance approaches. By aggregating rehabilitation intake across centers and standardizing clinical classifications, WildAlert helps quantify the broader extent of events that might otherwise remain unseen. Although rehabilitation records, carcass counts, and beach/stranding surveys each have inherent biases, combining these data streams can offer a more comprehensive, synergistic understanding of population-level impacts revealing patterns that extend far beyond the “tip of the iceberg.”

WildAlert’s success depends on the continued engagement of wildlife rehabilitation centers and the sustainability of data-sharing pipelines. The current implementation in five U.S. states already spans diverse ecosystems and operational contexts, but further expansion will require dedicated support, interoperability standards, and capacity-building in under-resourced regions. Nevertheless, the platform’s demonstrated scalability and real-time functionality position it as a valuable complement to existing passive and active surveillance systems, including those focused on zoonotic spillover, biodiversity loss, and climate-linked disease emergence.

WildAlert 2.0 represents a promising step toward operationalizing artificial intelligence for wildlife health surveillance. By integrating clinical data streams with contextual modeling and unsupervised anomaly detection, the system enables timely and scalable monitoring of emerging threats. As global environmental changes accelerate, the utility of integrated, adaptive platforms increases. WildAlert provides a model for how machine learning tools can be responsibly and effectively applied to enhance early warning capabilities, guide investigations, and ultimately inform on threats at the intersection of wildlife, ecosystem, and public health.

## Materials and Methods

### Establishment of the WildAlert early warning system

WildAlert was developed as a real-time, data-driven early warning system that utilizes case records submitted by wildlife rehabilitation organizations through the Wildlife Rehabilitation Medical Database (wrmd.org). At the time of writing, the system was deployed in Florida, California, Arizona, Pennsylvania and Washington, incorporating contributions from 182 rehabilitation organizations that provided continuous, real-time data streams (Fig 1). Data were processed through a machine learning model pipeline designed to perform syndromic tagging of cases and to detect statistical anomalies in admissions patterns across a range of wildlife taxonomic groups and clinical presentations (Fig. 1).

### Standardizing clinical classifications and circumstances of admission for wildlife rescue cases

Two natural language processing models were developed using the WRMD dataset. The first model assigned standardized clinical classifications to wildlife rescue cases based solely on free-text entries recorded at the time of admission, specifically reason(s) for admission, initial physical examination findings, and preliminary diagnoses. The second model classified cases according to circumstances of admission, also using free-text descriptions. The training dataset for clinical classifications was annotated by a board-certified wildlife veterinarian (T.K.), who assigned one or more clinical classifications to each case as appropriate. The dataset comprised 15,636 labeled observations across eleven possible clinical classifications (see Supplementary Materials), with 11,243 cases assigned a single classification and the remaining observations receiving multiple labels, including eight cases that were assigned five classifications. The circumstances of admission dataset comprised 35,844 cases annotated with 74 possible labels. This was annotated by experienced wildlife rehabilitators (D.D, R.A.) and a board-certified wildlife veterinarian (T.K.).

To ensure robust model evaluation, the datasets were randomly split into three subsets: training, validation, and testing. First, 15% of the data was reserved as an independent test set. The remaining 85% was further divided, with 20% allocated to validation and the remainder to training. We trained a custom large language model using Bidirectional Encoder Representations from Transformers (BERT) fine-tuned for multi-class, multi-label classification. Specifically, text data were processed using a pretrained DistilBERT tokenizer (*distilbert-base-uncased*), producing a sequence of token embeddings for each case note. Input text sequences were truncated or padded to a maximum length of 128 tokens. DistilBERT, a smaller and faster variant of BERT optimized for efficiency (Sanh et al., 2019), was selected to balance performance and computational cost. The large language models outperform conventional NLP models in bringing context of formal language structures and adapting to specialized domain vocabulary. Fine-tuning enabled the model to perform multi-label classification, allowing assignment of multiple pre-diagnostic clinical categories to a single case, capturing the complexity of wildlife rehabilitation, where animals often present with multiple clinical signs and/or co-occurring health conditions.

Model training was conducted using 10-fold cross-validation on the training datasets to optimize hyperparameters, including learning rate (2e-5), batch size (16), and number of epochs (5). A grid search strategy was employed to tune these parameters based on validation loss and F1-score. Class weights were applied to address moderate class imbalance, particularly for underrepresented labels such as Ocular Disease and Urogenital Disease. The best model was selected based on the highest macro-averaged F1-score on the validation set. Performance was evaluated on the independent dataset using accuracy, precision, recall, F1-score, and ROC AUC.

#### Hybrid anomaly detection using seasonal decomposition with autoencoder and isolation forest

To detect temporal anomalies across combinations of species, clinical classifications, and circumstances of admission, we developed a hybrid computational framework that integrates classical seasonal-trend decomposition with two complementary machine learning approaches: a neural network-based autoencoder and a tree-based Isolation Forest. This dual approach was designed to enhance the robustness and sensitivity of anomaly detection by leveraging both deep learning and ensemble-based techniques, each applied to the residual component of the time series after removal of systematic variation. For each taxon-classification combination, weekly admissions time series were decomposed into trend, seasonal, and residual components using an additive model, allowing isolation of long-term trends and recurring seasonal patterns. The residual component, representing non-systematic fluctuations, was retained as the primary signal for anomaly detection.

### Autoencoder-based anomaly detection

In the first approach, we employed a feedforward autoencoder trained on sliding windows of the residual time series. Each input window comprised *w* consecutive residual values, denoted as *x*_*i*_ = [*r*_*i*_, *r*_*i*+1_ …, *r*_*i*+*w*-_]. The autoencoder architecture consisted of an input layer of size *w*, followed by a hidden encoding layer with 64 neurons and *ReLU* activation, a bottleneck layer of dimension *d*, a decoding layer with 64 neurons, and an output layer of size *w* with linear activation. The model was optimized using the Adam optimizer and trained to minimize the mean squared error (MSE) between the input and reconstructed sequences over a fixed number of epochs. Following training, the model was used to reconstruct each residual window, producing a reconstruction 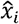.The reconstruction error for each window was computed as: 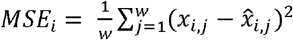. An anomaly threshold Ø was defined as the 95^th^ percentile of the distribution of the reconstruction errors and number of cases at a time point was flagged as anomalous if the reconstruction error exceeded the threshold and the final residual in the window was positive, i.e. *MSE*_*i*_ > ø and *r*_*i*+*w*-1_ > 0.

### Isolation forest-based anomaly detection

In parallel, we implemented a second anomaly detection strategy using the Isolation Forest algorithm. The residual values were reshaped into a one-dimensional feature space and used to train an Isolation Forest model with 100 estimators and a user-defined contamination parameter. The model assigned a binary anomaly label *α*_*i*_ ∈ {1, 1} and a continuous decision score *s*_*i*_ ∈ *R* to each residual *r*_*i*_. Anomalous high weekly case counts were defined as residuals that were both flagged as anomalous and had greater than zero. This hybrid framework enables sensitive and interpretable detection of biologically meaningful deviations in time series data, accommodating both gradual and abrupt changes in signal dynamics.

### Performance of anomaly detection methods

To compare the performance of anomaly detection methods, we computed a suite of agreement metrics between the Isolation Forest and Autoencoder outputs. These included Cohen’s Kappa for pairwise agreement, Jaccard similarity to quantify the proportion of overlapping anomaly events, and Fleiss’ Kappa to assess overall agreement across all three methods, including a standard deviation threshold-based rule. Additionally, we recorded the total number of anomalies detected by each method and the number of events jointly flagged by both machine learning models. To assess the robustness and comparative sensitivity of the models in the presence of gradual non-stationary behavior, we systematically evaluated their performance across a range of temporal and statistical conditions. A simulation framework was developed to test the effects of window size, contamination level, outlier injection, and slow-varying trends on anomaly detection outcomes. Two types of perturbations were introduced to mimic realistic challenges in longitudinal surveillance data: (i) stochastic outliers, injected as sporadic high-count spikes at random timestamps, and (ii) a gradual linear trend, modeled as an additive low-magnitude slope scaled to a 10% fraction of the maximum observed case count. This synthetic trend was injected using a deterministic function. 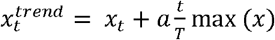, where *α* is a trend strength parameter (0.1), *t* is the timepoint, and *T* the total duration (number of observations) in the specific species clinical classification series. We iteratively evaluated both anomaly detection models across a range of sliding window sizes (3–20 weeks) and contamination levels (1–20%) with and without injected outliers and trends. For each parameter combination, we computed the number of anomalies detected by each model and the intersection (overlap) of detected events. Results were summarized separately for the 15 sporadic and 17 endemic WildAlert surveillance targets to evaluate how reporting patterns influence model sensitivity and stability. Ordinary least squares (OLS) regressions were fitted separately for endemic and sporadic datasets to quantify the relative influence of simulation parameters (outlier presence, trend addition, window size, and contamination level) on the number of detected anomalies. The resulting coefficients and explanatory power (R^2^) provided a standardized basis for comparing model robustness across the two reporting regimes.

### Assessing the timeliness and ecological sensitivity of WildAlert

To evaluate the timeliness and ecological relevance of WildAlert, we conducted a validation study using harmful algal bloom (HAB) events in Florida as natural experiments. Specifically, we focused on blooms of *Karenia brevis*, a dinoflagellate species known to produce neurotoxins (brevotoxins) that impact marine wildlife. We examined the temporal relationships of *K. brevis* cell counts and marine bird strandings using cross-correlation analysis. Spatiotemporally associated *K. brevis* cell counts with stranding locations were sourced from state monitoring programs. Marine birds tagged with either Neurological Disease or Nutritional Disease clinical classifications from Florida were selected as targets for exploration.

To evaluate the ecological sensitivity of WildAlert anomaly signals, we conducted a retrospective comparison against USDA-confirmed detections of highly pathogenic avian influenza (HPAI) in wild birds (https://www.aphis.usda.gov/livestock-poultry-disease/avian/avian-influenza/hpai-detections/wild-birds) using surveillance data collected by the California Department of Fish and Wildlife, the Florida Fish and Wildlife Conservation Commission, and USDA Wildlife Services and confirmed by the USDA National Veterinary Services Laboratories. Confirmatory detection periods were extracted from USDA surveillance records and overlaid onto WildAlert anomaly timelines for the corresponding species groups and states. For each surveillance target, we compared AE and IOF intensity scores (reconstruction error produced by a feedforward autoencoder trained on sliding windows of residual) during USDA-confirmed periods versus outside those intervals using Welch’s two-sample t-test and a non-parametric permutation test (1,000 iterations), allowing quantitative assessment of whether WildAlert signals were significantly elevated during periods of clustered HPAI detections, thereby providing an independent validation of anomaly responsiveness.

To evaluate lagged relationships between environmental variables and wildlife health outcomes, we conducted a cross-correlation analysis between weekly bird admissions and *K. brevis* cell counts, a proxy for harmful algal bloom (HAB) intensity. Bird admissions were treated as the dependent outcome. We computed Pearson’s cross-correlation coefficients *R*_*xy*_ across ±2, 4, 6, 8, and 10-week lags to identify temporal offsets that may indicate predictive relationships. Confidence intervals were calculated using a Bartlett-adjusted variance approach that accounts for autocorrelation in both time series (Bartlett, 1946). The Bartlett method computes the variance of the cross-correlation function using the product of autocorrelation functions of each series, providing a more conservative estimate of statistical significance. The analysis was performed on nine coastal bird species commonly affected by HABs and other environmental stressors: Great Egret (*Ardea alba*), Ruddy Turnstone (*Arenaria interpres*), Striated Heron (*Butorides striata*), Sanderling (*Calidris alba*), Tricolored Heron (*Egretta tricolor*), White Ibis (*Eudocimus albus*), Ring-billed Gull (*Larus delawarensis*), Black-crowned Night Heron (*Nycticorax nycticorax*), and Sooty Tern (*Onychoprion fuscatus*).

In addition to quantitative validation through cross-correlation analysis, we conducted a qualitative assessment of WildAlert’s real-world performance by reviewing wildlife health investigations that were triggered by system-generated alerts. Between June 2024 and January 2025, WildAlert flagged multiple anomalous events across Florida and California, prompting investigations and diagnostic follow-up by state wildlife agencies and wildlife rehabilitation organizations.

## Acknowledgments

The work was supported by Florida Fish and Wildlife Conservation Commission and the United States Fish and Wildlife Zoonotic Disease Initiative Grant Program award # P2380017. We thank all of the participating wildlife rehabilitation organizations in WildAlert across California, Florida, Arizona, Washington and Pennsylvania.

## Data Availability

Code available to train natural language processing models for various WildAlert tags are presented in respective GitHub repositories

Clinical Classification: https://github.com/PanditPranav/WildAlertModels

Circumstances for admission: https://github.com/PanditPranav/WildAlertModels_Circumstances

Code for anomaly detection models, their sensitivity analyses, and retrospective comparison with known events are presented in GitHub repository: https://github.com/PanditPranav/WildAlert_AnomalyModels. Wildlife rehabilitation data used in the manuscript is available on WildAlert: https://wildalert.org/ and can be accessed and visualized using the WildAlert platform.

