## Supplementary for "WildAlert: A Real-Time, AI-Driven Early Warning System for Wildlife Health and Ecological Threat Detection"

*Pranav S. Pandit.

**Supplementary Table 1.** Per-class precision, recall, and F1-score for the clinical classification model.

The model demonstrates high predictive performance across all 11 clinical categories derived from wildlife rehabilitation medical records.

| Clinical Classification | Precision | Recall | F1-score |
| --- | --- | --- | --- |
| Clinically Healthy | 0.97 | 0.97 | 0.97 |
| Dermatologic Disease | 0.94 | 0.88 | 0.91 |
| Gastrointestinal Disease | 0.95 | 0.89 | 0.91 |
| Hematologic Disease | 0.94 | 0.91 | 0.93 |
| Neurologic Disease | 0.94 | 0.9 | 0.92 |
| Nonspecific | 0.9 | 0.86 | 0.88 |
| Nutritional Disease | 0.94 | 0.92 | 0.93 |
| Ocular Disease | 0.92 | 0.89 | 0.9 |
| Physical Injury | 0.95 | 0.94 | 0.94 |
| Respiratory Disease | 0.92 | 0.9 | 0.91 |
| Urogenital Disease | 0.9 | 0.9 | 0.9 |

**Supplementary Table 2.** Per-class precision, recall, F1-score for the circumstances of admission model.

The model demonstrates high predictive performance across all 74 circumstances of admission derived from wildlife rehabilitation medical records.

| Circumstances of Admission | Precision | Recall | F1-score |
| --- | --- | --- | --- |
| Abduction with intent of rescue | 0.988 | 0.968 | 0.978 |
| Animal interaction | 0.965 | 0.928 | 0.946 |
| Bicycle collision | 0.984 | 0.968 | 0.976 |
| Born in captivity | 0.987 | 0.905 | 0.944 |
| Bow and Arrow | 1 | 1 | 1 |
| Cat interaction | 0.983 | 0.995 | 0.989 |
| Collision | 0.931 | 0.76 | 0.837 |
| Confiscation | 1 | 0.958 | 0.979 |
| Cooking oil exposure | 0.879 | 0.961 | 0.918 |
| Displaced from nest | 0.959 | 0.919 | 0.939 |
| Disturbed metabolic rest | 1 | 0.992 | 0.996 |
| Dog interaction | 0.985 | 0.99 | 0.988 |
| Domestic animal interaction | 0.985 | 0.99 | 0.988 |
| Dumped | 0.969 | 0.854 | 0.908 |
| Electrocution | 1 | 0.175 | 0.299 |
| Entrapment | 0.955 | 0.653 | 0.775 |
| Entrapped in building | 0.919 | 0.86 | 0.889 |
| Entrapped in chimney | 0.956 | 0.632 | 0.761 |
| Entrapped in fence | 0.969 | 0.98 | 0.975 |
| Entrapped in fishing tackle | 0.985 | 0.98 | 0.983 |
| Entrapped in litter / garbage | 0.976 | 0.974 | 0.975 |
| Entrapped in netting / string / wire | 0.95 | 0.976 | 0.963 |
| Entrapped in storm drain / sewer | 0.992 | 0.992 | 0.992 |
| Entrapped in vehicle | 0.969 | 0.979 | 0.974 |
| Entrapped in water | 0.95 | 0.959 | 0.954 |
| Fire / smoke | 0.992 | 0.969 | 0.981 |
| Friendly | 0.984 | 0.956 | 0.97 |
| Garden / farm equipment collision | 0.996 | 0.989 | 0.993 |
| Grounded | 0.924 | 0.88 | 0.901 |
| Gunshot | 0.992 | 0.993 | 0.993 |
| Hand held object collision | 1 | 0.91 | 0.953 |
| Illness | 0.864 | 0.717 | 0.784 |
| Inappropriate human intervention | 0.992 | 0.98 | 0.986 |
| Maladaptation / failure to thrive | 0.982 | 0.82 | 0.893 |
| Mating injury | 1 | 0.854 | 0.921 |
| Nest / habitat disturbance or destruction | 0.951 | 0.976 | 0.963 |
| Non-domestic animal interaction | 0.965 | 0.954 | 0.959 |
| Non-weapon projectile | 0.936 | 0.82 | 0.874 |
| Nuisance animal | 0.95 | 0.959 | 0.955 |
| Orphan | 0.95 | 0.969 | 0.96 |
| Paint exposure | 1 | 0.74 | 0.851 |
| Pet | 0.962 | 0.93 | 0.945 |
| Petrochemical exposure | 0.987 | 0.899 | 0.941 |
| Physical trauma | 0.906 | 0.915 | 0.91 |
| Plane collision | 1 | 1 | 1 |
| Poisoned | 0.988 | 0.983 | 0.986 |
| Powerline / wire collision | 0.941 | 0.957 | 0.949 |
| Referral / transfer | 0.965 | 0.652 | 0.778 |
| Same species interaction | 0.957 | 0.929 | 0.942 |
| Stranded | 0.957 | 0.943 | 0.95 |
| Surrender | 0.971 | 0.957 | 0.964 |
| Tar exposure | 0.954 | 0.939 | 0.947 |
| Toxic exposure | 0.958 | 0.958 | 0.958 |
| Train collision | 0.987 | 0.991 | 0.989 |
| Trapped in glue trap | 0.996 | 0.99 | 0.993 |
| Trapped in humane / cage trap | 0.938 | 0.789 | 0.857 |
| Trapped in leghold / trap / snare | 0.967 | 0.981 | 0.974 |
| Tree trimming | 0.989 | 0.989 | 0.989 |
| Unauthorized or untrained rehabilitation | 0.927 | 0.85 | 0.887 |
| Undetermined | 0.94 | 0.903 | 0.921 |
| Vehicle collision | 0.982 | 0.99 | 0.986 |
| Watercraft collision | 1 | 0.333 | 0.5 |
| Weather event | 0.979 | 0.986 | 0.983 |
| Wind turbine collision | 0.967 | 0.956 | 0.961 |
| Window / wall collision | 0.986 | 0.991 | 0.988 |

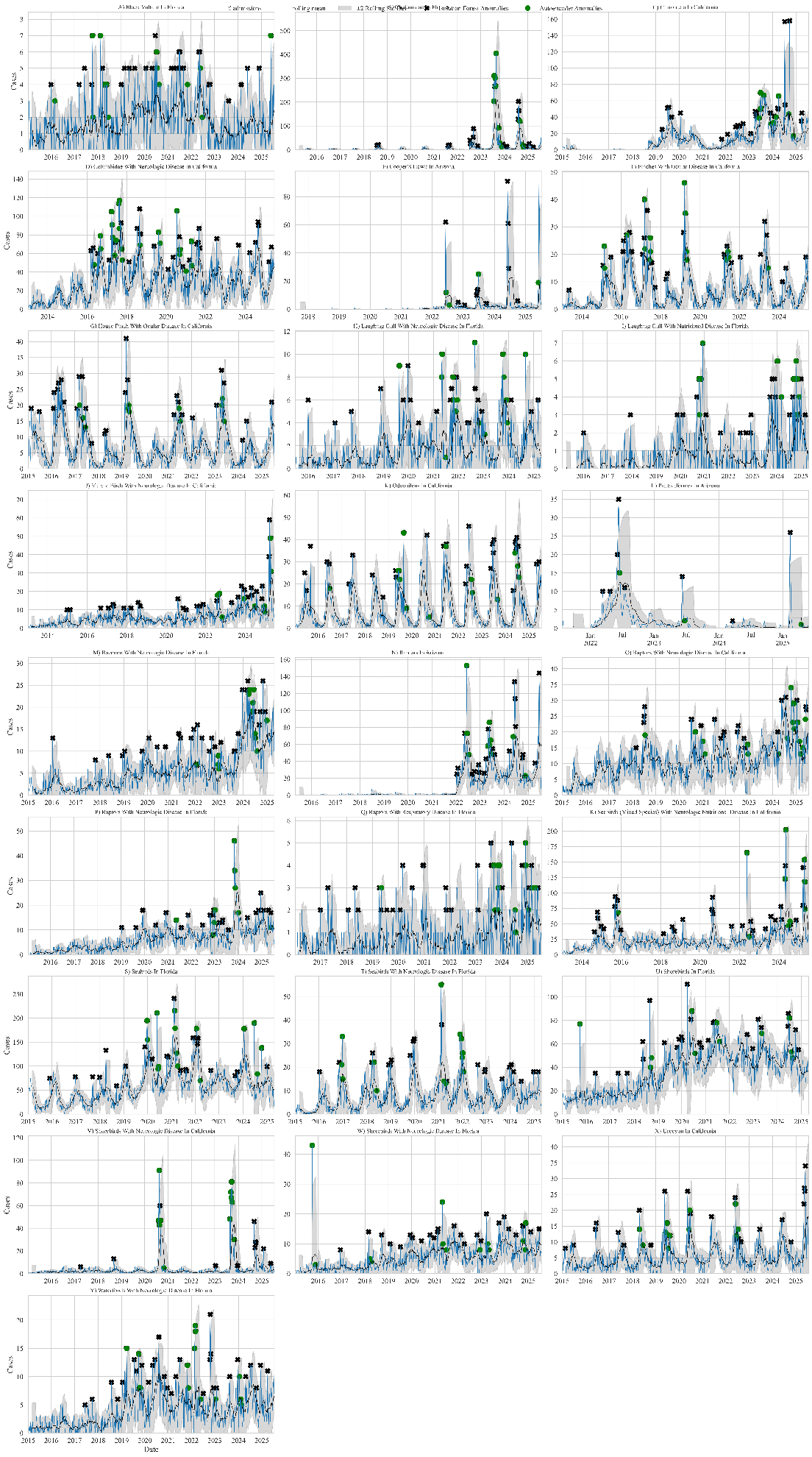

Supplementary Fig 1: WildAlert surveillance target time series for tracking various health events in wildlife across commonly found species and clinical classification. Blue lines represent monthly admissions, black lines indicate the rolling mean, and gray shaded areas show the rolling standard deviation. Anomalies detected by the Isolation Forest model are marked with black stars, and anomalies identified by the Autoencoder model are shown as green circles.

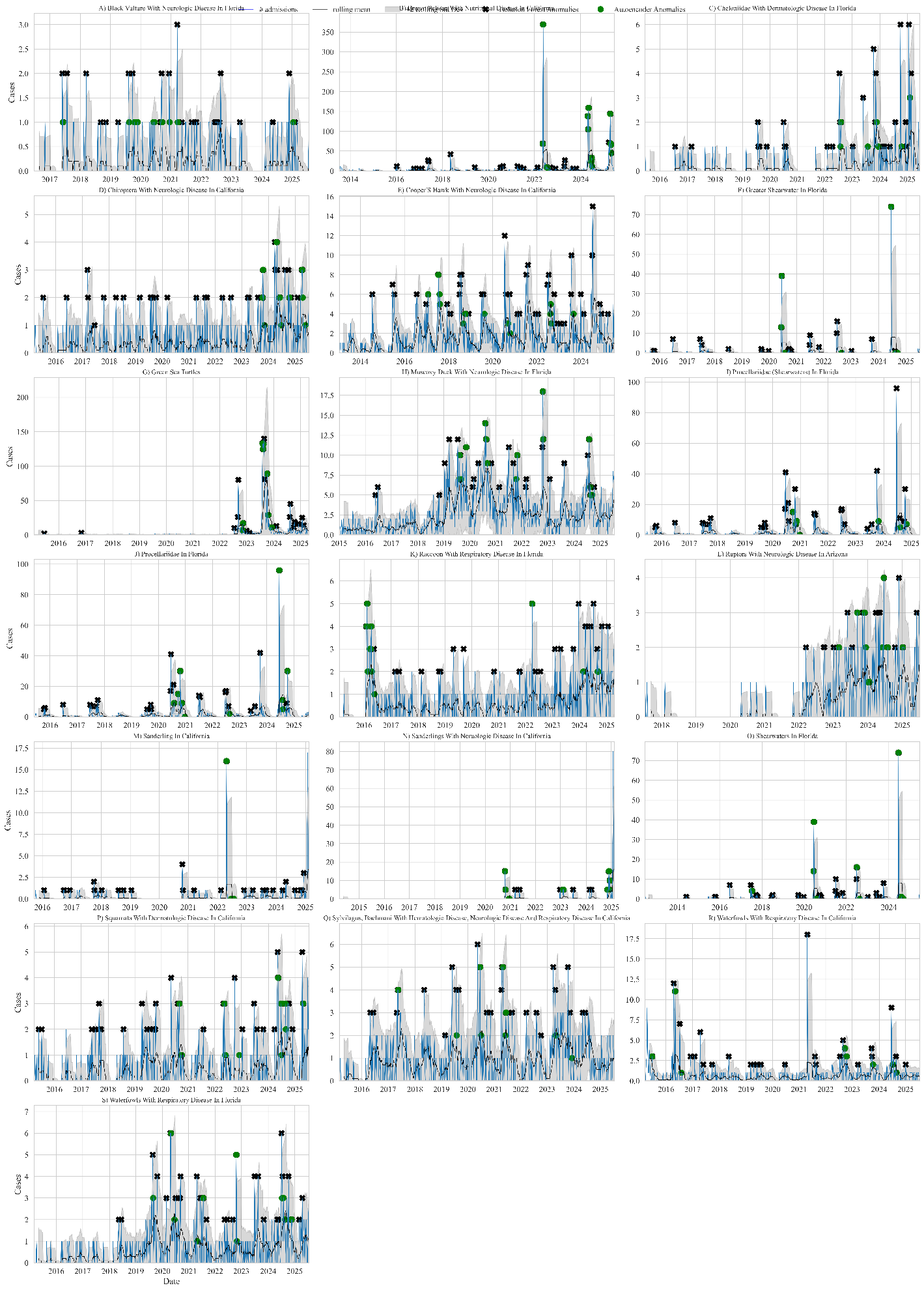

Supplementary Fig 2: WildAlert surveillance target time series for tracking various health events in wildlife across sporadic, rare combinations of species and clinical classification. Blue lines represent monthly admissions, black lines indicate the rolling mean, and gray shaded areas show the rolling standard deviation. Anomalies detected by the Isolation Forest model are marked with black stars, and anomalies identified by the Autoencoder model are shown as green circles.
